# Optical Nanosensors for Real-time Feedback on Insulin Secretion by β-Cells

**DOI:** 10.1101/2021.03.21.435763

**Authors:** Roni Ehrlich, Adi Hendler-Neumark, Verena Wulf, Dean Amir, Gili Bisker

## Abstract

Quantification of insulin is essential for diabetes research in general, and for the study of pancreatic β-cell function in particular. Herein, fluorescent single-walled carbon nanotubes (SWCNT) are used for the recognition and real-time quantification of insulin. Two approaches for rendering the SWCNT sensors for insulin are compared, using surface functionalization with either a natural insulin aptamer with known affinity to insulin, or a synthetic PEGylated-lipid (C_16_-PEG(2000Da)-Ceramide), both of which show a modulation of the emitted fluorescence in response to insulin. Although the PEGylated-lipid has no prior affinity to insulin, the response of C_16_-PEG(2000Da)-Ceramide-SWCNTs to insulin is more stable and reproducible compared to the insulin aptamer-SWCNTs. The C_16_-PEG(2000Da)-Ceramide-SWCNTs optical response is excitation-wavelength dependent, where resonant excitation leads to a larger fluorescence decrease in response to insulin. The SWCNT sensors successfully detect insulin secreted by β-cells within the complex environment of the conditioned media. The insulin is quantified by comparing the SWCNTs fluorescence response to a standard calibration curve, and the results are found to be in agreement with an enzyme-linked immunosorbent assay (ELISA). This novel analytical tool for real time quantification of insulin secreted by β-cells provides new opportunities for rapid assessment of β-cell function, with the ability to push forward many aspects of diabetes research.

## 1. Introduction

Diabetes mellitus is a group of metabolic diseases caused by impaired insulin secretion and defective insulin action, which lead to chronic hyperglycemia,^[1]^ effecting over 400 million people worldwide.^[2]^ Insulin is a 5.8 kDa naturally occurring peptide hormone which is synthesized and secreted by pancreatic β-cells.^[3]^ The main role of insulin is to preserve the blood glucose level by promoting cellular glucose uptake, as well as regulating the metabolism of lipids, proteins, and carbohydrates.^[4]^ The most potent stimulus of insulin secertion is glucose.^[5]^ An increase in blood glucose level induces β-cell electrical activity, which produces an elevated Ca^+2^ concentration that triggers exocytosis of insulin granules.^[6]^ Type 1 diabetes is characterized by autoimmune destruction of the β-cells, thus requiring exogenous insulin administration as therapy.^[1]^ Type 2 diabetes is characterized by β-cell dysfunction causing relative insulin deficiency and insulin resistance.^[7]^ In order to understand the pathogenesis of diabetes and the mechanisms involved in the deterioration of β-cells, considerable effort has been made in understanding the physiological processes which lead to insulin production and secretion in β-cells.^[8]^ Many investigations have attempted to understand the mechanism of insulin secretion when triggered by a rise in the extracellular glucose concentration.^[9]^ Understanding these mechanisms can lead to the development of new therapies for diabetes and contribute to the possibility of engineering insulin producing cells for cell replacement therapys.^[10,11]^

There is a greatly restricted supply of viable human pancreatic islets which limits the opportunity for studies of β-cell function.^[12]^ Many attempts have been made over time to establish successful insulin-secreting β-cell lines which retain normal regulation of insulin secretion.^[11]^ One the most common insulin-secreting β-cell lines are βTC cells.^[11]^ βTC lines are derived from insulinomas in transgenic mice expressing SV40 T antigen (Tag) and show normal glucose regulated insulin secretion.^[13]^ βTC-tet cells are derived from the same type of mice, only in which the SV40 T antigen is under control of the tetracycline gene regulatory system. By shutting off Tag expression in βTC-tet cells in the presence of tetracycline, growth arrest can be induced, leading to a gradual increase in their insulin content while maintaining normal insulin production and secretion.^[14]^ In recent years, there have been some advancements in creating human pancreatic β-cell lines which secrete insulin in response to glucose, with the hope of pushing forward the study of β-cell biology and drug discovery.^[15–17]^

The ability of β-cells to produce, store and release insulin is crucial for defining β-cell function.^[18]^ There is an increasing demand for simple insulin detection methods which would benefit clinical diagnostics as well as research.^[19]^ The main analytical methods for insulin quantification are immunoassays such as enzyme-linked immunoassays (ELISA),^[20]^ radioimmunoassay (RIA),^[21]^ chemiluminescence immunoassay (CLIA),^[22]^ and chromatography methods.^[19]^ ELISA can detect a target antigen in a sample through the color change obtained by using an enzyme-linked conjugate and an enzyme substrate.^[23]^ Sandwich type ELISA are the most common assays for the detection of insulin, having a low limit of detection in the pM range.^[19]^ However, ELISA requires multiple incubation and washing steps, such that the duration of the assay is usually over two hours. RIAs, which were the first widely used methods for the detection of insulin,^[24,25]^ consist of labeling the antigen or the antibody by a radioactive isotope.^[26]^ For insulin detection, unlabeled insulin samples and radiolabeled antigen compete for a limited number of antibodies, and the ratio between the bound and unbound antigen is used for insulin quantification. Due to safety concerns regarding the radiolabeled antigen, this method has been gradually replaced by other options.^[19]^ In CLIAs, the label, which indicates the analytical reaction, are chemiluminescent molecules.^[27]^ In the case of insulin detection, an antibody coats the test plate, and a secondary antibody with a chemiluminescent label, changes its intensity in the presence of insulin, which can then be correlated to the insulin concentration. CLIAs have high signal intensity as well as shorter incubation times, but have a relatively high cost.^[19]^ Therefore, a new method for a rapid, low cost, and simple insulin detection would highly benefit the research of β-cell function.

Recently, an insulin sensor based on single walled carbon nanotubes (SWCNTs) functionalized by lipid-poly(ethylene glycol) (PEG) was discovered.^[28]^ The unique optical and electronic properties of SWCNTs make them favorable as fluorescent sensors for biomedical applications,^[29–35]^ due to their fluorescence being in near-infrared (nIR) range, where biological samples are mostly transparent.^[36,37]^ Further, they do not show photobleaching nor blinking,^[38]^ and were shown to be biocompatible long term *in vivo*.^[39–41]^ Functionalized SWCNTs can detect targets of interest using a heteropolymer that is adsorbed onto the SWCNTs surface via non covalent interactions such that it recognizes a specific target analyte.^[42,43]^ The binding of the target molecule modifies the spectral properties of the nIR fluorescence emission of the SWCNTs by either intensity changes or wavelength shifts,^[30,44]^ both of which can be optically detected. This method was successfully applied to detect riboflavin, L-thyroxine, oestradiol,^[43]^ neurotransmitters,^[45]^ nitroaromatics,^[46,47]^ NO,^[48]^ H_2_0_2_,[49–51] small volatile molecules and odors,^[52,53]^ lipid,^[54]^ as well as larger macromolecules such as the protein fibrinogen.^[42]^ Recently, a screening of PEGylated lipids revealed a corona phase-SWCNT complex, C_16_-PEG(2000Da)-Ceramide-SWCNT, that can be used as an optical sensor for insulin,^[28]^ showing a florescence intensity decrease in the presence of insulin. Insulin was recognized and quantified by the SWCNT sensors both in buffer and in the serum environment.^[28]^

A different approach for SWCNT-based molecular recognition relies on biomolercular binding elements such as antibodies,^[55–57]^ aptamers,^[58,59]^ and other binding partners.^[60–70]^ Aptamers are short, single stranded nucleic acids which are selected for their specific target using the SELEX procedure (systematic evolution of ligands by exponential enrichment).^[71]^ Often, their high affinity to the targets results from a conformational effect as they may fold around the target upon binding.^[72]^ An natural aptamer for insulin was found within the insulin gene promoter, and was shown to be able to capture human insulin from standard solutions as well as from nuclear extracts of pancreatic cells.^[73]^ An optical response to insulin using SWCNTs can be achieved by anchoring the insulin aptamer onto the surface of the SWCNT, by combining a DNA anchor sequence such as (AT)_15_ with the aptamer binding domain.^[58]^

In this work, we compare and contrast both approaches for SWCNT-based insulin recognition, namely the synthetic PEGylated-lipid and the natural aptamer functionalization, and find the PEG-lipid-SWCNT response to be more stable and reproducible. Further, we find that the fluorescence response of the PEG-lipid-SWCNT insulin sensor depends on the nanotube chirality and that resonant excitation is benifical in terms of the extent of response. Finally, we demonstrate the detection and quantification of insulin secreted by βTC-tet pancreatic cells. **(Figure 1)**. Our result pave the way to a simple, real time, sensing and quantification of insulin secreted by β-cells which could greatly contribute to the study of β-cell function and diabetes research.

**Figure 1.**
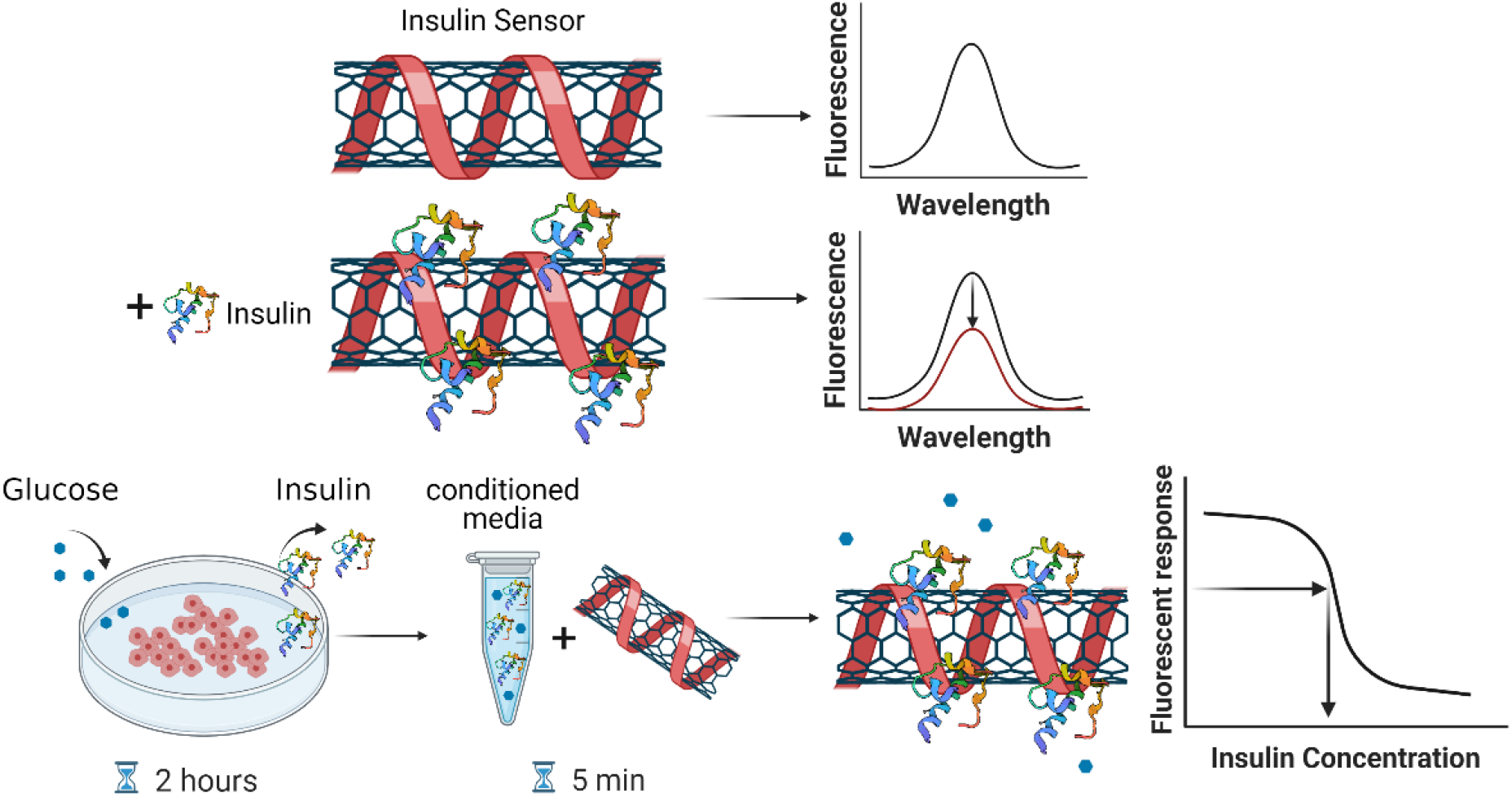
Schematic illustration of the SWCNT insulin sensor. The functionalized SWCNT fluoresce in the nIR, where insulin binding results in a modulation of the emitted fluorescence. For the insulin secretion assay, glucose is added to insulin-secreting β-cells for an incubation time of 2 hours. Subsequently, the conditioned media is collected and added to the SWCNT sensors solution, and the recorded optical response following a short incubation of 5 minutes is used to infer the concentration of the secreted insulin. Insulin illustration was adapted from protein data base (PDB) entry 1ZEH.^[74]^ Created with BioRender.com

## 2. Results

### 2.1 SWCNT sensors comparison

In order to render the SWCNT sensors for insulin, soudium cholate (SC) suspended SWCNTs were dialyzed with either C_16_-PEG(2000Da)-Ceramide^[28]^ or (AT)_15_-insulin aptamer^[58]^ to exchange the SC wrapping. The resulting suspensions showed clear fluorescence emission peaks in the nIR and a significant florescence response with the addition of insulin. In the presence of 33 µg ml^-1^ insulin, the C_16_-PEG(2000Da)-Ceramide**-**SWNCTS showed a 29% decrease in florescence intensity of the (10,2) chirality **(Figure 2a)**, and the (AT)_15_-Insulin Aptamer-SWCNTs showed an intensity decrease of 32% of the (6,5) chirality, as well as a 16% decrease and 10 nm wavelength redshift of the (10,2) chirality **(Figure 2b)**.

**Figure 2.**
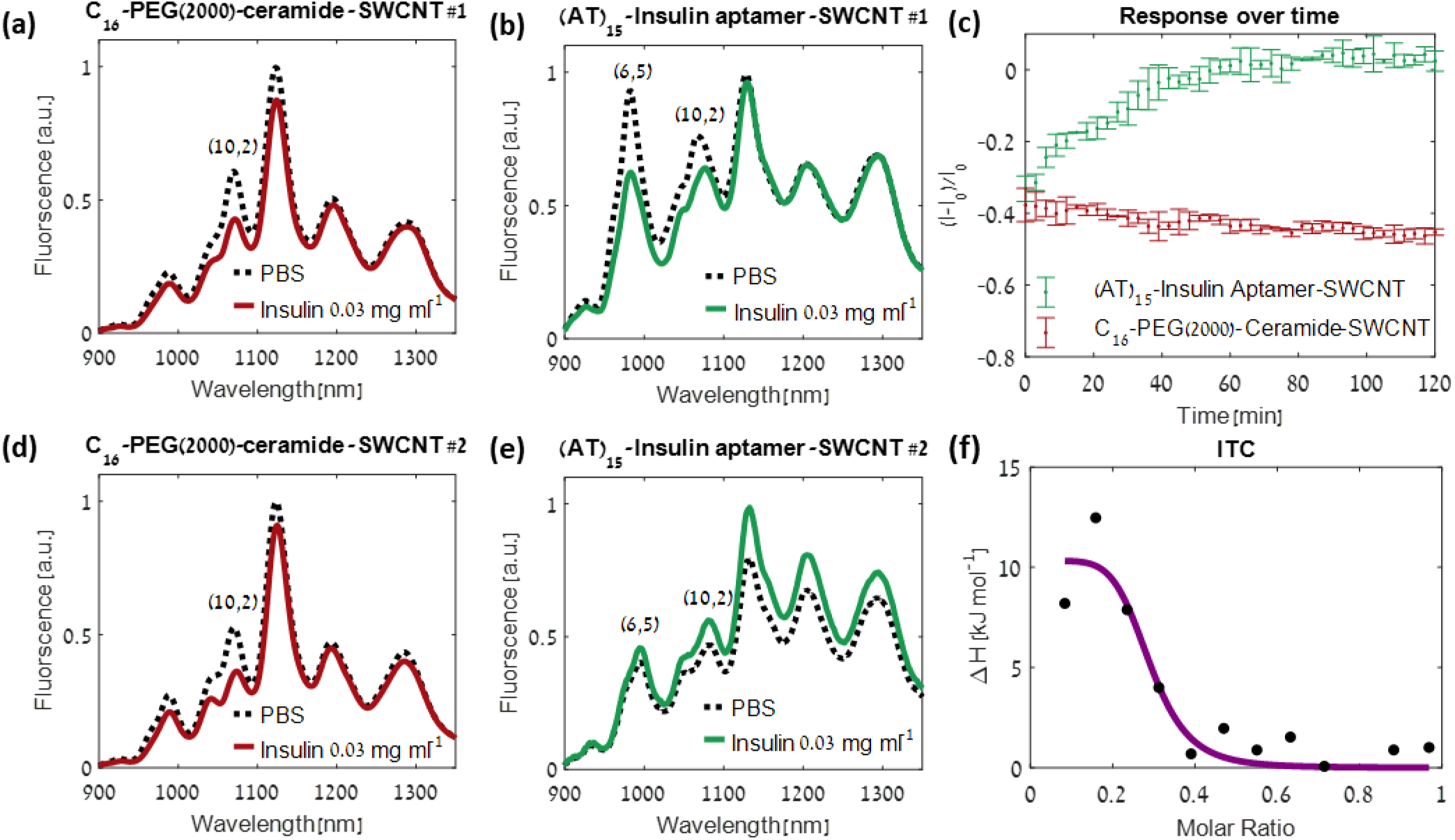
(a) Fluorescence emission of C_16_-PEG(2000Da)-Ceramide-SWCNT (1 mg L^−1^) sensor following the addition of insulin (33 μg ml^−1^, solid red curve), or following the addition of an equal volume of phosphate buffered saline (PBS) as control (dashed black curve) (b) Fluorescence response of (AT)_15_-Insulin Aptamer-SWCNT (1 mg L^−1^) sensor to insulin (33 μg ml^−1^, solid green curve), or following the addition of an equal volume of PBS as control (dashed black curve). (c) Relative fluorescence response over the duration of two hours of C_16_-PEG(2000Da)-Ceramide-SWCNT and (AT)_15_-Insulin Aptamer-SWCNT after the addition of insulin (0.06 mg ml^−1^). The C_16_-PEG(2000Da)-Ceramide-SWCNT response remains stable over time and the (AT)_15_-Insulin Aptamer-SWCNT response diminishes over time. The relative responses was calculated for the chirality with the highest response, (10,2) for C_16_-PEG(2000Da)-Ceramide-SWCNT and (6,5) for (AT)_15_-Insulin Aptamer-SWCNT. (d) Fluorescence response of batch #2 of C_16_-PEG(2000Da)-Ceramide-SWCNT (1 mg L^−1^) sensor to insulin (33 μg ml^-1^, solid red curve), or following the addition of an equal volume of PBS as control (dashed black curve), showing a reproducible response to insulin. (e) Fluorescent response of batch #2 of (AT)_15_-Insulin Aptamer-SWCNT (1 mg L^−1^) sensor to insulin (33 μg ml^−1^,solid green curve), or following the addition of an equal volume of PBS as control (dashed black curve), showing opposite responses to insulin. (f) ITC thermogram for the titration of insulin into insulin-aptamer solution. The curve indicates an endothermic binding reaction.

The response of the two functionalized SWCNT insulin sensors were compared by measuring the fluorescence response over time after the addition of insulin to SWCNT suspensions. While the C_16_-PEG(2000Da)-Ceramide**-**SWNCTS showed a stable fluorescence response over a duration of two hours, the response of the (AT)_15_-Insulin Aptamer-SWCNTs diminished over time, and the initial fluorescence was recovered after ∼40 minutes (**Figure 2c** and **Figure S1**). Comparing different batches, the dialysis of SC-SWCNTs with C_16_-PEG(2000Da)-Ceramide showed a reproducible response to insulin, **(Figure 2a and 2d)**, whereas the surfactant exchange from SC-SWCNTs to (AT)_15_-Insulin Aptamer via dialysis showed a batch to batch variation with opposite response trends **(Figure 2b and 2e)**. Interestingly, according to Isothermal titration calorimetry (ITC), C_16_-PEG(2000Da)-Ceramide has no prior affinity to insulin.^[28]^ In contrast, ITC measurements confirmed the affinity between the insulin aptamer and insulin **(Figure 2f)**. Nevertheless, the C_16_-PEG(2000Da)-Ceramide SWCNTs showed a stable and reproducible optical response to insulin. Owing to its sensing performance, the C_16_-PEG(2000Da)-Ceramide SWCNT was chosen for further investigation, over the (AT)_15_-Insulin Aptamer-SWCNT.

### 2.2 C_16_-PEG(2000Da)-Ceramide SWCNTs characterization

The C_16_-PEG(2000Da)-Ceramide-SWNCTS showed a bright fluorescent emission under 500 - 840 nm laser excitations as seen in the excitation-emission profile, where each peak corresponds to a different SWCNT chirality **(Figure 3a)**. In order to further investigate the sensor response to insulin, the excitation emission profile was recorded with the presence of 0.06 mg ml^-1^ Insulin. The normalized fluorescence response (Δ*I*/*I*_0_) interestingly revealed excitation-wavelength dependent response **(Figure 3b)**, where resonant excitation in the E_11_ transitions^[75]^ led to larger extent of fluorescence intensity decrease. Of the chiralities which showed the largest response to insulin, we chose to focus on the (10,2) chirality due to its brightness and the ease of ditingushing its emission from neighboring peaks.

**Figure 3.**
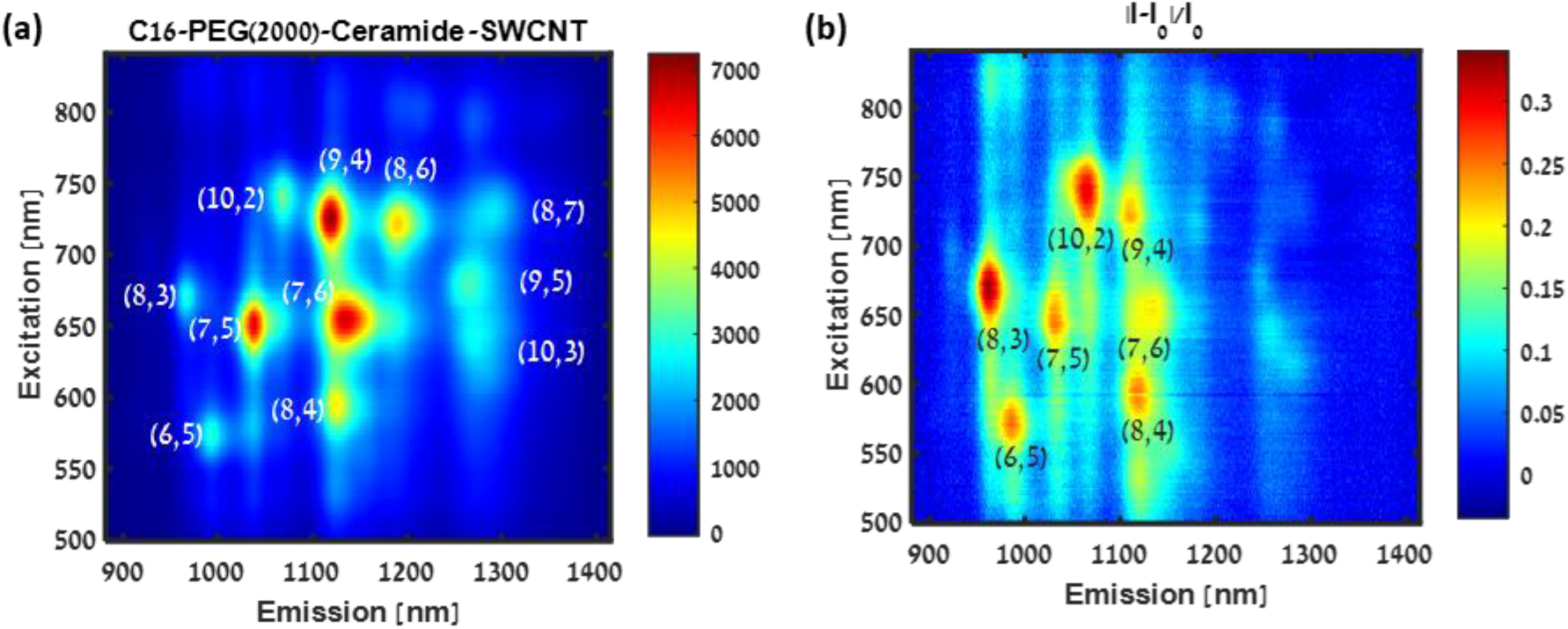
(a) Excitation–emission profile of the C_16_-PEG(2000Da)-Ceramide-SWCNT (1 mg L^−1^) sensor (b) Excitation–emission profile of the relative fluorescence response (Δ***I***/***I***_***0***_) of C_16_-PEG(2000Da)-Ceramide-SWCNT (1 mg L^−1^) sensor to insulin (0.06 mg ml^−1^), showing a chirality and excitation-wavelength dependent response.

The fluorescence spectra of the C_16_-PEG(2000Da)-Ceramide-SWNCTs were then recorded with increasing concentrations of insulin in PBS, showing a gradual decrease in fluorescence as previously reported^[28]^ **(Figure 4a)**, and deconvoluted to the individual contributions of the different SWCNT chiralities. To further emphasize the importance of resonant excitation, a variety a wavelengths ranging from 670 nm to 785 nm, including the 742 nm excitation resonance of the (10,2) chirality, were chosen for excitation. The data was fitted using the Hill isothermal model^[76]^

**Figure 4.**
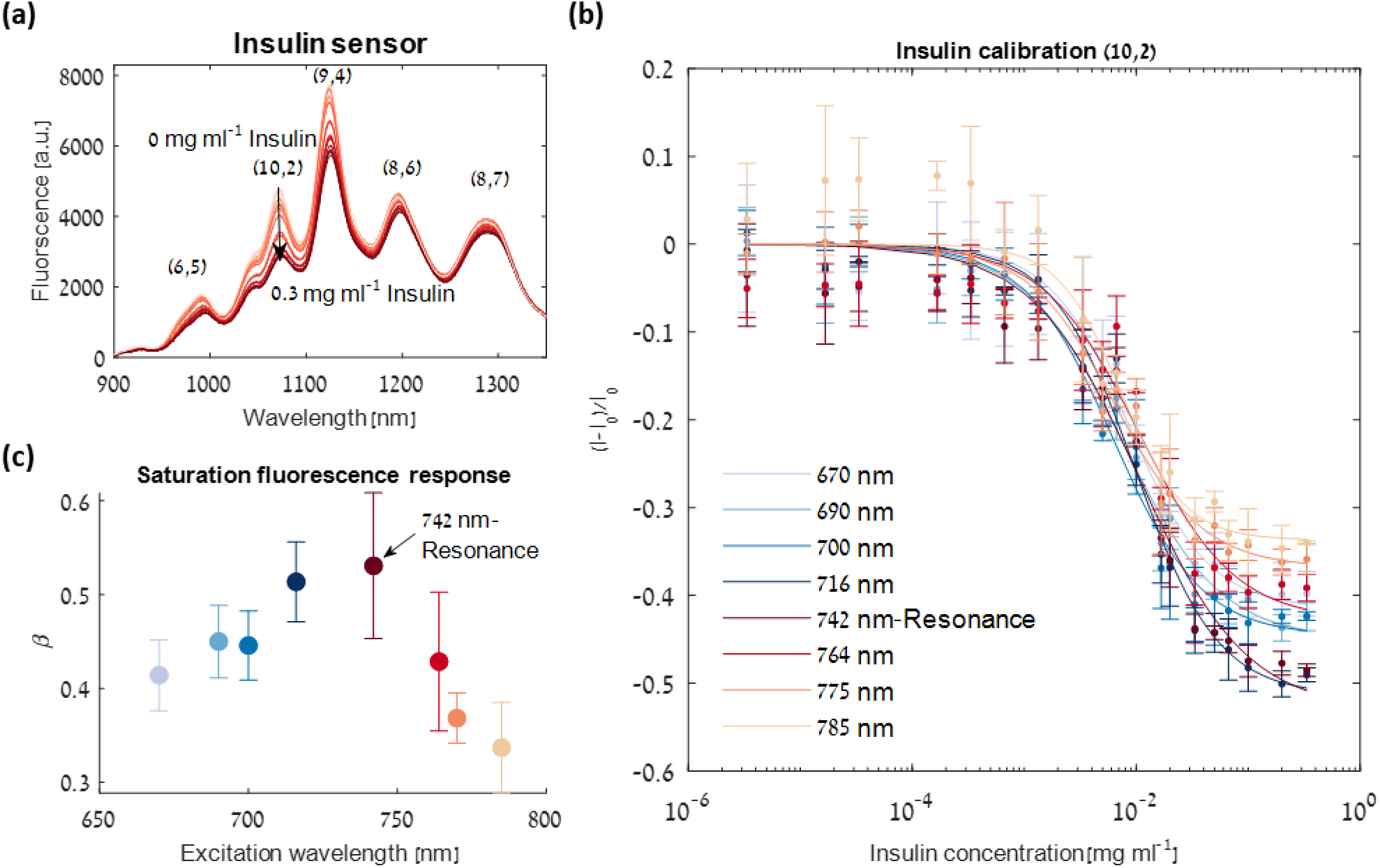
(a) Fluorescence emission spectra of C_16_-PEG(2000Da)-Ceramide-SWCNTs under 742 nm laser excitation (resonance for (10,2) chirality) with 0, 3.3 ×10^−6^, 1.6 ×10^−5^, 3.3 ×10^−5^, 1.6 ×10^−4^, 3 × 10^−4^, 6× 10^−4^, 1 × 10^−3^, 3 × 10^−3^, 1.6 × 10^−3^, 5 × 10^−3^, 6 × 10^−3^, 1 × 10^−2^, 1.6 × 10^−2^, 2 × 10^−2^, 3× 10^−2^, 5 × 10^−2^, 6× 10^−2^, 0.1, 0.2 and 0.3 mg ml^−1^ insulin show a gradual decrease in emission intensity with increasing insulin concentration. (b) Calibration curves of C_16_-PEG(2000Da)-Ceramide-SWCNTs versus insulin concentration at the peak emission wavelength of the (10,2) chirality under various excitation wavelengths. The solid lines represent the fit according to equation (1). A maximal response at saturation is received when exciting at 742 nm, the (10,2) excitation resonance. (c) The β parameter and its 95% confidence intervals versus the various excitation wavelengths. This fit parameter quantifies the maximal saturation response showing maximal response for resonant excitation.

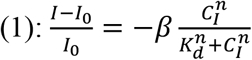

where *I*_0_ is the initial fluorescence intensity, *I* is the final fluorescence intensity, *β* is a proportionality factor and the maximal relative response at saturation, *C_I_* is the insulin concentration, *K_d_* is the dissociation constant, and *n* is the Hill coefficient (**Table S1**). Plotting the different calibration curves for the (10,2) chirality at its peak emission wavelength and different excitation wavelengths **(Figure 4b)** shows a maximal response at saturation when exciting at the (10,2) excitation resonance, 742 nm, as quantified by the *β* parameter **(Figure 4c**). We thus conclude that 742 nm resonant excitation provides optimal optical conditions for insulin sensing and quantification.

### 2.3 Insulin Secretion assay

In order to demonstrate the applicability of the insulin sensor for rapid quantification of insulin in a complex environment, we challenged the C_16_-PEG(2000Da)-Ceramide**-**SWNCTS in a β-cells insulin secretion assay.

Initially, the fluorescent intensity modulation of the C_16_-PEG(2000Da)-Ceramide**-**SWNCTS in response to a variety of insulin concentrations was measured in Krebs-Ringer HEPES buffer (KRHB), to rule out any possible effect of the cell media on the sensor performance compared to PBS. The data were fitted using the hill isothermal model,^[76]^ providing a calibration curve for the optical response, and a limit of detection of 0.13 µg ml^-1^ insulin (**Figure 5a**, red line). The sensor response in KRHB was comparable to its response in PBS, as quantifies by the three fit parameters and their 95% confidence intervals **(Table S2)**, confirming there is no significant response to any component of the medium.

**Figure 5.**
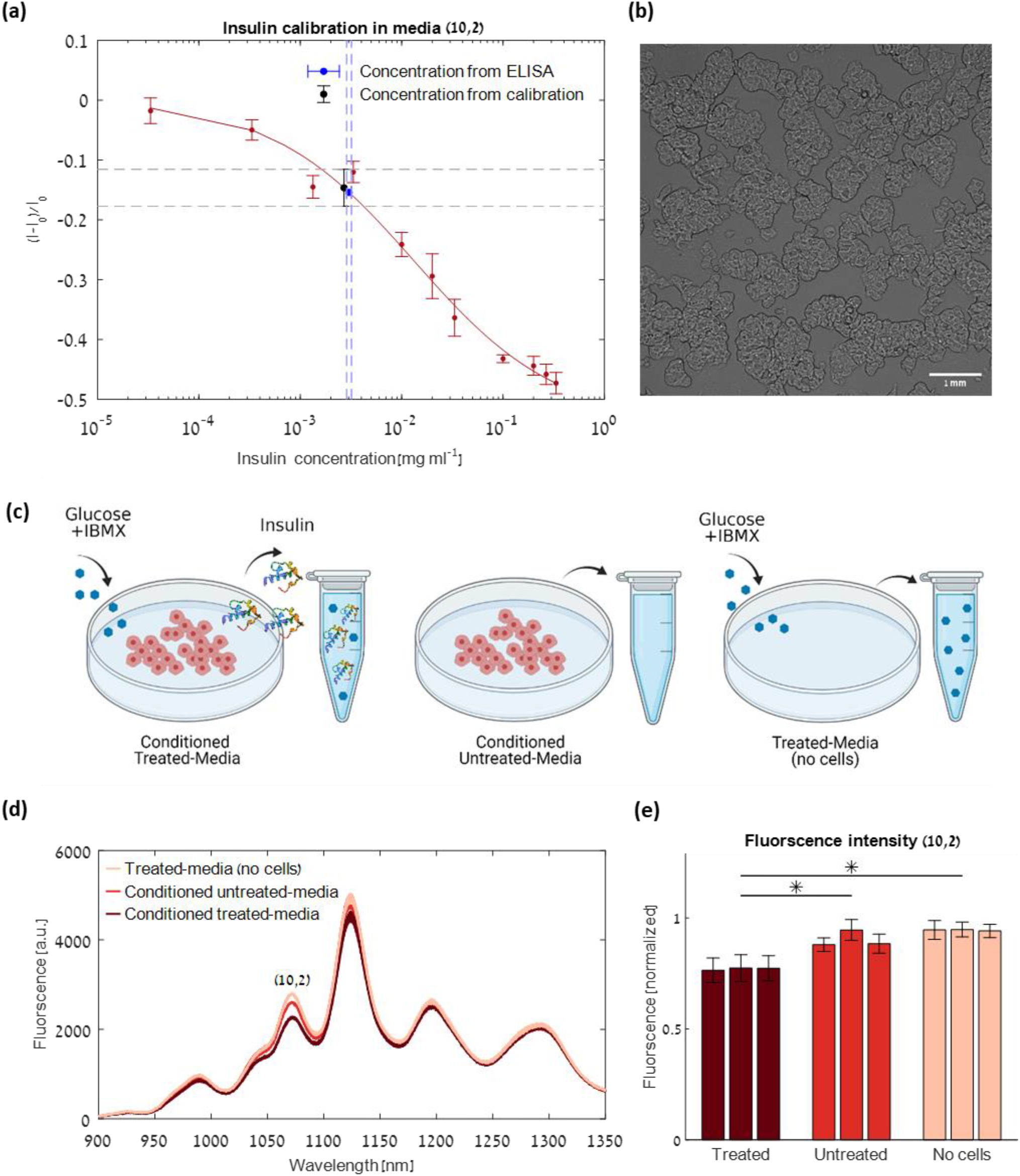
(a) Calibration curve of C_16_-PEG(2000Da)-Ceramide-SWCNTs in KRHB media versus insulin concentration for peak emission wavelength of the (10,2) chirality (red dots). The solid line represents the fit according to the model described in the text. The blue dot represents the insulin concentration according to ELISA in the conditioned treated media, with the corresponding STD (dashed blue lines). The black dot represents the normalized fluorescence response in the conditioned treated media of the C_16_-PEG(2000Da)-Ceramide-SWCNT sensor compared to the conditioned untreated media, with the corresponding STD (dashed gray lines). The calibration curve falls within the cross section of the two measurement value ranges, demonstrating a successful insulin quantification using the SWCNT sensor. (b) Bright-field image of the βTC-tet cells used in the insulin secretion assay. (c) The three conditions used for the insulin secretion assay: 1. Conditioned KRHB media from βTC-tet cells treated with 16 mM glucose and 0.5 mM IBMX for induced insulin secretion 2. Control: Conditioned KRHB media from untreated βTC-tet cells and 3. Reference: KRHB media treated with 16 mM glucose and 0.5 mM IBMX (no cells) (d) Fluorescence emission spectra of C_16_-PEG(2000Da)-Ceramide-SWCNTs in the three conditions. A 4.6% intensity decrease of the (10,2) chirality was observed between the control and the reference and a 14% intensity decrease between the control and the conditioned treated media. (e) Normalized (10,2) chirality fluorescence of the C_16_-PEG(2000Da)-Ceramide-SWCNT sensor for the triplicates of the three different conditions, showing fluoresce intensity decrease in the treated samples as a result of the secreted insulin. Error bars represent the STD between three replicate measurements. Statistical analysis was done using the Mann-Whitney U-test.^[81]^ *p-value<0.001

For the secretion assay, insulin secreting βTC-tet cells **(Figure 5b)** were incubated for two hours with KRHB containing 16 mM glucose and 0.5 mM IBMX to stimulate insulin secretion.^[14]^ Cell cultures that were not treated with glucose\IBMX were used as a control, and cells-free samples of KRHB containing 16 mM glucose and 0.5 mM IBMX were used as a reference **(Figure 5c)**.

Samples of the treated and untreated conditioned media as well as the reference samples of KRHB with 16 mM glucose and 0.5 mM IBMX in the absence of cells were diluted by a factor of 3 with C_16_-PEG(2000Da)-Ceramide**-**SWNCTs and incubated for a short period of 5 min, and their fluorescence spectra were recorded. The fluorescence intensity showed a 14±3% intensity decrease of the (10,2) chirality in the treated conditioned media compared to the untreated conditioned media, indicating the successful detection of the secreted insulin by the SWNCT insulin sensor **(Figure 5d and 5e)**. According to this normalized fluorescence response of the C_16_-PEG(2000Da)-Ceramide**-**SWNCTs insulin sensor and the calibration curve, the concentration of the secreted insulin in the treated conditioned-media was evaluated to be 8.07±0.15 µg ml^-1^ (Figure 5a, gray dashed lines). Given the initial cells density and the duration of incubation prior to the secretion assay, we estimated to have had ∼6× 10^8^ cells at the beginning of the glucose-stimulus experiment, for which this secreted insulin concentration falls within the expected range.^[14]^ Comparing to the reference sample (media + glucose\IBMX only, no cells), the untreated conditioned media led to a 4.6% intensity decrease which was attributed to the general cells secretome,^[77–80]^ whereas the treated conditioned media led to an 18% fluorescence intensity decrease, which stems from the combined effect of both the secreted insulin and other components of the secretome. In order to rule out the possibility of an optical response resulting from IBMX or glucose, the C_16_-PEG(2000Da)-Ceramide**-**SWNCTS were tested in the presence of 0.5 mM IBMX or 16 mM glucose, showing no change in the fluorescent signal **(Figure S2)**.

The gold standard analytical technique for protein quantification is an ELISA,^[19]^ which utilizes antibodies for the recognition and fluorescent labels for quantification. In order to validate our results, the concentration of insulin secreted by the cells was determined by ELISA. The secreted insulin concentration in the conditioned media of the βTC-tet cells treated with glucose and IBMX was found to be 9.1±0.59 µg ml^-1^ (Figure 5a, blue dashed lines), which is in very good agreement with the calculated 8.07±0.15 µg ml^-1^ concentration according to the fluorescence response of our SWCNT sensor. The untreated cells retained a basal insulin secretion of 0.069±0.027 µg ml^-1^, which is below the limit of detection of the SWCNT sensor, confirming that the fluorescence response of these samples resulted from other secretome factors within the conditioned media, and thus should be taken as the baseline for comparing the fluorescence emission of the treated conditioned media, as we have done.

## 3. Discussion

The study of β-cell function is crucial for the advancement of diabetes research^[11,82–84]^. Since the key role of β-cells is to efficiently store and secrete insulin, there is an increasing demand for novel methods for rapidly quantifying insulin secretion from these cells.

We engineered two nanosensors for insulin utilizing either C_16_-PEG(2000Da)-Ceramide or insulin aptamer functionalization of fluorescent SWCNTs, both of which showed a modulation of the fluorescence emission upon the interaction with insulin. Although ITC measurement confirmed the binding affinity between insulin and the insulin aptamer (Figure 2f) and ruled out any affinity between insulin and the C_16_-PEG(2000Da)-Ceramide,^[28]^ we found the synthetic PEG-lipid-SWCNTs to be preferable in terms of stability and reproducibility of the optical response (Figure 2a, 2b, 2c, 2d and 2e).

The florescence intensity of the C_16_-PEG(2000Da)-Ceramide-SWNCTs immediately decreased upon the exposure to insulin, and was shown to be stable for a duration of two hours (Figure 2c). This result manifests the key role played by the specific conformation adopted by the PEG-lipid or aptamer when adsorbed onto the nanotube surface, which can greatly affect the sensors response.^[42,43,85]^ Taken together, the C_16_-PEG(2000Da)-Ceramide SWCNT was chosen for further experimentation for insulin detection and quantification. The fluorescence modulation of the C_16_-PEG(2000Da)-Ceramide SWCNTs interestingly showed an excitation wavelength dependent response to insulin (Figure 3b), where resonant excitation led to a larger fluorescence decrease. Specifically, we focused on the (10,2) chirality peak emission, whose contribution to the fluorescence can be easily distiguished from neighboring peaks. By scanning a variety of excitation wavelengths, we demonstrated that the excitation of the (10,2) chirality at its resonance excitation wavelength (742 nm) resulted in the largest relative fluorescence response at saturation (Figure 4c).

As a proof of concept, we used a pancreatic β-cell line for glucose-induced insulin secretion assay and demonstrated the detection and quantification of insulin in the conditioned media following the glucose stimulus. The calculated insulin concentration according to the calibration curve of our SWCNT sensor, was in an excellent agreement with ELISA (Figure 5a), which is the gold-standard today for quantification of proteins. ^[19]^

Insulin ELISA kits contain multiple-well plates coated with insulin antibodies, and the assay requires many incubation steps with different reagents (such as the enzyme conjugate, the enzyme substrate, washing buffer, blocking buffer, and a stopping solution), resulting in an exceedingly long process ranging from two hours up to five hours or more. At the end of the process, the absorption is measured at a specific wavelength and compared to known concentration references for quantification. Usually, insulin ELISA kits have a detection limit in the ng ml^-1^ range.

In contrast to ELISA, our C_16_-PEG(2000Da)-Ceramide-SWNCTs insulin sensor provides a rapid feedback on insulin concentration following a short incubation of 5 minutes. Further, it is 100% synthetic thus having high chemical and thermal stability, and can be stored in 4°C rather than -20°C, there is no need for any reagents or extra solutions besides the SWCNT sensors and the sample, and no special sample preparation is required. The main challenge of the SWCNT sensor is extending its detection limit which was calculated to be 0.13 µg ml^-1^, and it will be the subject of future research.

## 4. Conclusion

In conclusion, we have compared two different approaches for insulin quantification using functionalized SWNCT, and found that C_16_-PEG(2000Da)-Ceramide-SWCNT to be a stable and reproducible insulin nanosensor with optical signal transduction. We showed that the fluorescence response is dependent on the excitation wavelength and demonstrated the importance of resonant excitation for optimal sensor performance in terms of the response at saturation. Our sensor can successfully detect and quantify insulin secreted by pancreatic β-cells with only a short incubation time of 5 minutes, providing real time feedback. This work paves the way to a simple analytical tool for the quantification of insulin which could accelerate β-cell research. Owing to the SWCNT fluorescence in near-infrared range, where biological samples are mostly transparent^[36]^, this methodology can be extended for insulin detection in additional settings that can benefit from rapid feedback including, for example, within the microenvironment of intact pancreatic islets,^[86]^ or in the proximity of an insulin injection site.^[87–89]^

## 5. Methods

### 5.1 SWCNTs suspension

HiPCO SWCNTs (NanoIntegris) were suspended in 2 wt% SC (Sigma-Aldrich), using bath sonication (80 Hz for 10 minutes, Elma P-30H), followed by direct tip sonication (12 W for 60 minutes, QSonica Q125). The suspension was then ultracentrifuged (41,300 rpm for 4 hours, OPTIMA XPN – 80) in order to separate the individually suspended SWCNTs from aggregates and impurities.^[28]^ Subsequently, 40 mg L^-1^ of SC-SWCNTs were mixed with 2 mg ml^-1^ C_16_-PEG(2000Da)-Ceramide (Avanti Polar Lipids) or 1 mg ml^-1^ (AT)_15_-Insulin Aptamer^[58]^ (ATATATATATATATATATATATATATATAT GGT GGT GGG GGG GGT TGG TAG GGT GTC TTC) (Integrated DNA Technologies) and dialyzed against water for 5 days with multiple water exchanges. In this process, the PEGylated lipid or ssDNA absorb onto the SWCNTs thus exchanging the SC wrapping.^[28]^ The absorption spectra of the suspensions were recorded using an ultraviolet-visible-nIR (UV-Vis-nIR) spectrophotometer (Shimadzu UV-3600 PLUS), where sharp distinguishable peaks indicated a successful suspension **(Figure S3)**.

### 5.2 Sensor characterization

For characterizing sensor response, 1 mg L^-1^ of the suspended SWNCTs were added to the wells of a 96 well plate to which insulin (Sigma-Aldrich) was added at a final concentration of 33 µg ml^-1^. The fluorescence spectra were acquired using a nIR microscope coupled to a liquid-nitrogen cooled InGaAs detector, using a spectrograph (PyLoN-IR 1024-1.7 and HRS-300SS, Princeton Instruments, Teledyne Technologies).

For excitation-emission maps, the excitation wavelengths were tuned using a super-continuum white-light laser (NKT-photonics, Super-K Extreme) coupled to a tunable bandpass filter (NKT-photonics, Super-K varia, Δλ = 20 nm).

For insulin calibration curves 1 mg L^-1^ SWNCT-C_16_-PEG(2000Da)-Ceramide was added to the wells of a 96 wells plate to which insulin was added at final concentrations ranging from 3.3 ng ml^-1^ to 0.3 mg ml^-1^. Following 30 min of incubation, the fluorescence spectra were acquired.

### 5.3 Cell culture

β-tc tet cells were incubated at 37°C, 5% CO_2_, and cultured in DMEM medium containing 25 mM glucose and supplemented with 10 % Fetal Bovine Serum (FBS), 1% penicillin-streptomycin, and 1% L-glutamine (complete DMEM). The cells were subcultured at about 90% confluency using 0.25% Trypsin solution containing 0.05% EDTA (all purchased from Biological Industries).

### 5.4 Insulin secretion assay

β-tc tet cells were seeded in a 100 mm cell culture dishe at a density of 4.8 × 10^6^ cells. When the cells were 80% confluent, the growth medium was supplemented with 1 μg ml^-1^ tetracycline (Sigma-Aldrich) for 7 days to induce growth arrest, which results in higher insulin content.^[14]^ On the day of the assay, cells were rinsed twice with KRHB (HEPES (10 mM) (Biological Industries), NaHCO_3_ (25 mM) (Daejung), NaH_2_PO_4_ (2 mM) (Sigma-Aldrich), MgSO_4_ (1 mM) (Carlo Erba), KCl (5 mM) (Sigma-Aldrich), CaCl_2_ (2.5 mM) (Daejung), NaCl (118 mM) (Bio-Lab Chemicals) and then pre-incubate in KRHB at 37°C for 1 h. The medium was then replaced with either fresh KRHB or KRHB containing IBMX (0.5 mM) (Sigma-Aldrich) and glucose (16 mM) (Millipore) and incubated for 2 hours. In addition, culture plates were incubated with KRHB containing IBMX (0.5 mM) and glucose (16 mM) with no cells to serve as a reference. Each condition was repeated in triplicates. Following the 2 h incubation, the medium was removed and centrifuged at 300 g for 3 min to remove any detached cells. The conditioned media samples, as well as the cell-free media control sample, were incubated for 5 minutes with 1 mg L^-1^ SWNCT-C16-PEG(2000Da)-Ceramide and the fluorescence spectra was acquired. Each sample was tested in triplicates. Moreover, the insulin concentration was determined by ELISA (Alpco Mouse insulin ELISA 80-INSMS-E01) following manufacturer”s instructions.

### 5.5 ITC

Isothermal titration calorimetry (MicroCal PEAQ-ITC) measurements were done by a series of 3 µL injections of insulin stock (50 µM) solution into the isothermal titration calorimetry cell containing insulin aptamer (10 µM) or PBS (Bio Prep) as a control.

## Acknowledgements

G. Bisker acknowledges the support of the Zuckerman STEM Leadership Program, the Israel Science Foundation (grant No. 456/18), The Ministry of Science, Technology, and Space, Israel (grant No. 3-17426), and the Nicholas and Elizabeth Slezak Super Center for Cardiac Research and Biomedical Engineering at Tel Aviv University. R. Ehrlich is supported by The Ministry of Science, Technology, and Space, Israel. D. Amir is supported by the Glaser Foundation and The Marian Gertner Institute for Medical Nanosystems. The authors thank Prof. Limor Landsman and Lina Sakhneny for valuble discussions.

