## Supporting Information for "Optical Nanosensors for Real-time Feedback on Insulin Secretion by β-Cells"

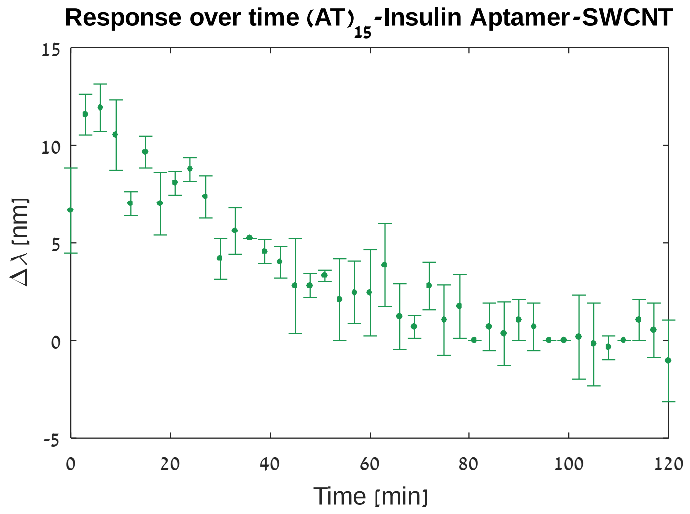


**Figure S1**: Response over the duration of two hours of $(AT)_{15}$-Insulin Aptamer-SWCNT to insulin (0.06 mg $ml^{-1}$). The observed red shift for the (6,5) chirality diminishes over time.

**Table S1:** Fit parameters and their 95% confidence intervals used to fit the data in figure 4b according to equation 1.

| $\lambda_{ex}$ [nm] | $\beta$ | $K_d$ [mg ml$^{-1}$] | $n$ |
|---|---|---|---|
| **670** | 0.414 (0.376, 0.451) | 0.008 (0.006, 0.011) | 1.214 (0.845, 1.582) |
| **690** | 0.449 (0.411, 0.488) | 0.007 (0.005, 0.009) | 0.994 (0.745, 1.242) |
| **700** | 0.445 (0.408, 0.482) | 0.006 (0.004, 0.008) | 1.104 (0.793, 1.415) |
| **716** | 0.513 (0.471, 0.556) | 0.009 (0.007, 0.012) | 1.174 (0.873, 1.474) |
| **742** | 0.530 (0.452, 0.608) | 0.010 (0.005, 0.015) | 0.905 (0.589, 1.221 |
| **764** | 0.428 (0.354, 0.502) | 0.010 (0.004, 0.016) | 1.014 (0.551, 1.476) |
| **775** | 0.368 (0.341, 0.395) | 0.006 (0.004, 0.008 | 1.105 (0.835, 1.375) |
| **785** | 0.336 (0.288, 0.385) | 0.007 (0.004, 0.010) | 1.531 (0.5911, 2.47) |

**Table S2:** Fit parameters and their 95% confidence bounds intervals obtained by fitting the data according to equation 1 in the presence of PBS or KRHB, $\lambda_{ex} = 742\ nm$. The sensor response in KRHB is comparable to its response in PBS, as quantified by the three fit parameters, indicating no significant response to any component of the buffer.

| Buffer | $\beta$ | $K_d$ [mg ml$^{-1}$] | $n$ |
|---|---|---|---|
| **KRHB** | 0.538 (0.438, 0.638) | 0.013 (0.002, 0.024) | 0.616 (0.400, 0.833) |
| **PBS** | 0.530 (0.452, 0.608) | 0.010 (0.005, 0.015) | 0.905 (0.589, 1.221) |

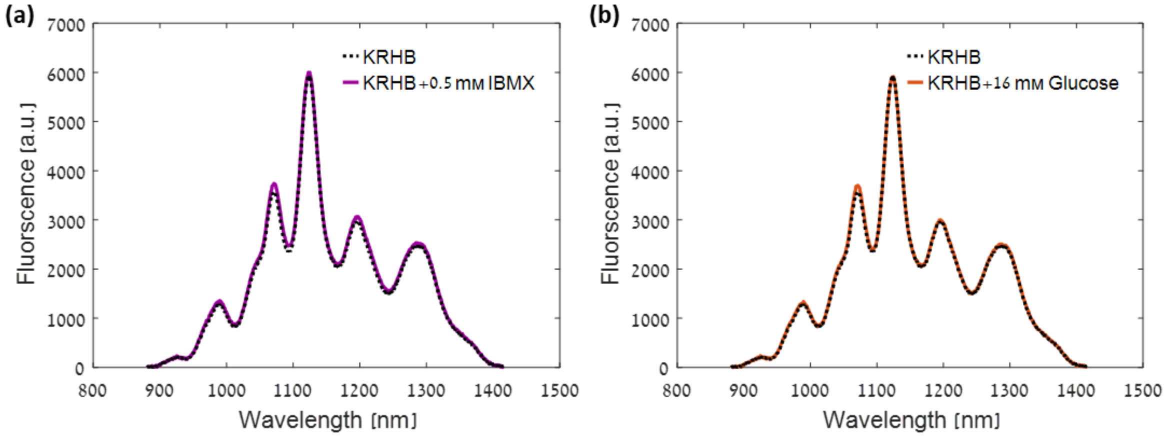


**Figure S2**: $C_{16}$-PEG(2000Da)-Ceramide-SWNCTS in the presence of IBMX (0.5 mM) or glucose (16 mM), show no change in the fluorescent signal.

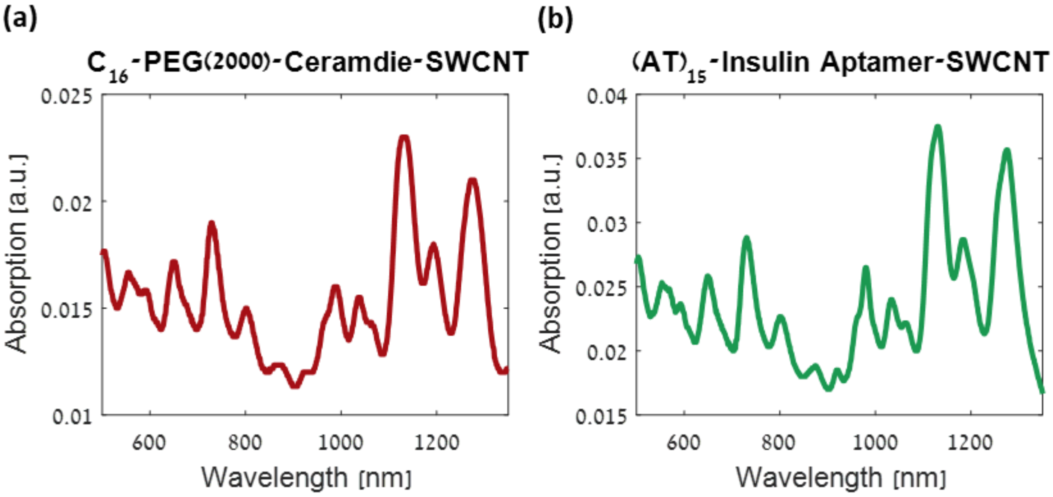


**Figure S3:** Absorption spectra of the two SWCNT suspensions after dialysis.
